# Intraspecific variation stabilizes classic predator-prey dynamics

**DOI:** 10.1101/2021.09.27.461947

**Authors:** Stefano Allesina, Zachary R. Miller, Carlos A. Serván

**Affiliations:** Department of Ecology & Evolution, University of Chicago Chicago, IL 60637; Northwestern Institute on Complex Systems, Northwestern University; Department of Mathematics, University of Chicago Chicago, IL 60637

**Keywords:** predator-prey, consumer-resource, intraspecific variation, phenotypic variation

## Abstract

In 1920, Alfred J. Lotka found that, to his “considerable surprise”, the dynamics of a simple predatorprey model he had devised led “to undamped, and hence indefinitely continued, oscillations” ^1,2^— which he thought epitomized the “rhythm of Nature” dear to the Victorians. In 1926, the same model was proposed independently by mathematician Vito Volterra ^3,4^, who was inspired by the work of his son-in-law, fish biologist Umberto D’Ancona ^5^. For over a century, the equations that now bear their names have served as a template for the development of sophisticated models for population dynamics ^6–10^. Coexistence in this classic predator-prey model is fragile—stochasticity or temporal variability in parameter values result in extinctions. The dynamics can be stabilized by intraspecific competition or other forms of self-regulation, but the prevalence of these processes in large food webs has been questioned ^11,12^. Here we show that when we consider populations characterized by intraspecific variability, dynamics are stable—despite the absence of any direct self-regulation. Our results can be generalized further, defining a new class of consumer-resource models ^8,13^. By accounting for intraspecific variation, which is manifest in all biological populations, we obtain dynamics that differ qualitatively and quantitatively from those found for homogeneous populations—challenging a central assumption of many ecological models.

---

The classic Lotka-Volterra predator-prey model describes the dynamics of a prey population, with population size or density *x*(*t*), and a predator population, *y*(*t*), in time. In its simplest formulation, the model can be written as:

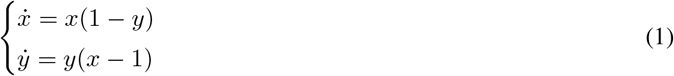

where 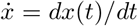 and 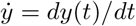 are the derivatives of the population sizes with respect to time. The model has two equilibria: a trivial one where both prey and predators are absent, and the coexistence equilibrium *x*\* = *y*\* = 1. As proved independently by Lotka ^1,2^ and Volterra ^3,4^, whenever the two populations are initialized at positive values, the dynamics lead to indefinite cycles around the coexistence equilibrium. In the phase plane, trajectories cycle counter-clockwise describing a curve determined by the initial conditions (Fig. 1). In fact, the coexistence equilibrium is neutrally stable, and one can identify a “constant of motion”, i.e., a quantity that is conserved through the dynamics ^2,14^. The same qualitative result is found when considering more general parameterizations (Methods).

Coexistence in this classic model is fragile: when, for example, we introduce stochasticity or allow parameters to vary in time, dynamics invariably and rapidly lead to extinction.

**Figure 1:**
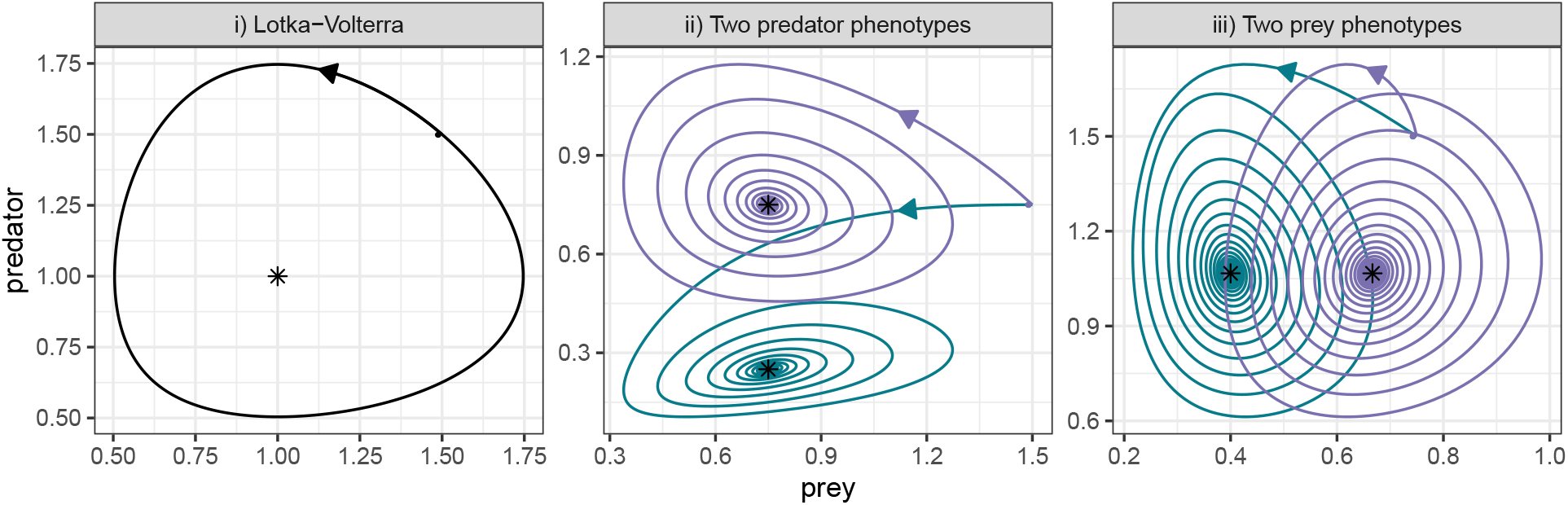
Global stability of predator-prey models incorporating intraspecific variation. i) Population dynamics for the classic Lotka-Volterra predator-prey model in Eq. 1. Dynamics (solid line) cycle counterclockwise around the coexistence equilibrium (asterisk), starting from any initial condition (solid point). In this model, populations are completely homogeneous. When we consider populations composed of two phenotypes linked by reproduction, dynamics are globally stable (Methods): ii) two predator phenotypes (colored trajectories), Eq. 2, *δ* = 1/2; iii) two prey phenotypes, Eq. 3, *ϵ* = 1/4.

This behavior is clearly incompatible with the widespread coexistence observed in natural systems. To account for it, the model can be extended, for example by introducing self-regulation (e.g., intraspecific competition, or other forms of density dependence) for the predator or prey populations. The modified model yields globally stable coexistence (i.e., attained irrespective of the positive initial conditions, Methods). These models can be scaled up to include an arbitrary number of predators and prey (e.g., as in MacArthur’s consumer-resource model ^8^), potentially exhibiting global stability when self-regulation is sufficiently strong ^13^.

The strength and prevalence of self-regulation is crucial for determining the outcomes in such models. Random-matrix approaches, following May’s pioneering work ^15^, show that predator-prey interactions tend to be stabilizing ^16^—all else being equal, ecological communities dominated by predator-prey interactions require weaker self-regulation to be stable than those characterized by other types of interaction. However, further extensions of these approaches show that even in food webs, stability requires self-regulation to be widespread, and quite substantial ^12^. Because many ecological models rely on self-regulation (usually arising from intraspecific competition) to produce stable dynamics, these processes play a central role that cuts across much of coexistence theory.

Here we demonstrate that intraspecific variation provides an alternative route to stabilization. We introduce variation in the predator or prey by considering populations composed of distinct phenotypes coupled by reproduction. For example, suppose that the predator population consists of two phenotypes (e.g., male and female ^17^) with characteristic mortality rates. By modifying Eq. 1, we obtain:

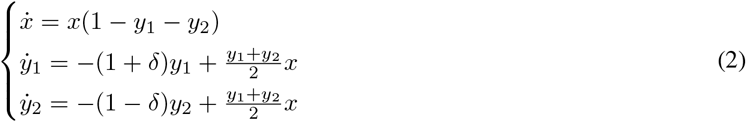

In Eq. 2, the prey suffers from both predator phenotypes equally, the first predator phenotype has elevated mortality (0 < *δ* < 1), and the second phenotype has reduced mortality. The two predator phenotypes pool their reproduction, with the term 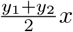 modeling the fact that, when reproducing, male and female individuals are produced with equal probability. For the special case *δ* = 0, the model reduces to Eq. 1. For any other value of *δ* (i.e., any difference between phenotypes), however, the dynamics are globally stable: the predator phenotypes and the prey settle at their respective equilibrium abundances from any positive initial condition (Fig 1, Methods).

Similarly, we can consider a model in which there are two prey phenotypes (e.g., vulnerable and defended), sharing reproduction and experiencing different attack rates from the predator:

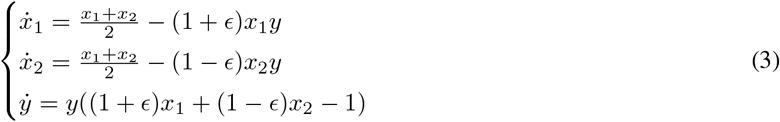

In Eq. 3, the shared reproduction of the prey phenotypes appears in the linear term, and for any value 0 < *ϵ* < 1 we find globally-stable dynamics, invariably leading to the coexistence equilibrium (Fig. 1, Methods).

These results can be generalized to an arbitrary number of phenotypes, modeling, for example, a distribution of mortality rates for the predator. In this case, as well, the coexistence equilibrium is always globally stable (Methods). As an example, Fig. 2 shows the dynamics for a system in which the prey has a single phenotype, while the predator has three (each characterized by a specific mortality rate), and offspring are assigned to each phenotype with a given probability.

**Figure 2:**
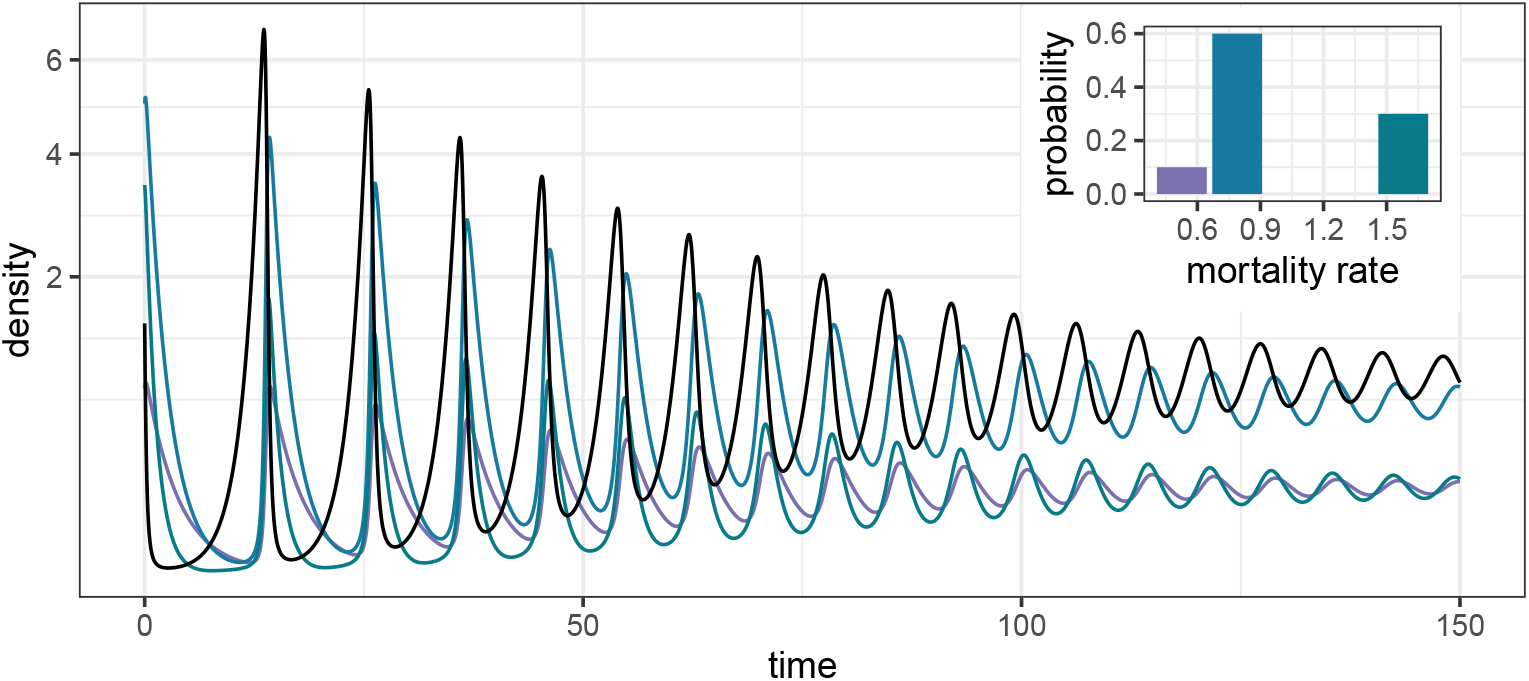
Modeling a distribution of mortality rates for the predator. Stable dynamics obtained when the prey has a single phenotype (black), while the predator has three (colors). Contrary to the form of Eq. 2, offspring are not assigned to each phenotype with equal probability (histogram in the inset). Dynamics are globally stable, as can be proven for an arbitrary number of predator phenotypes and distribution of mortality rates (Methods).

Stability arises in these models through damping of the predator-prey oscillations. As the predator (prey) population decreases, differential survival of the phenotypes results in a shift toward higher relative abundance of the lower-mortality type(s). When predator (prey) populations are increasing, offspring of the lower-mortality type(s) are “dissipated” into the more vulnerable phenotypic class(es). The net result is that the effective trait value for the multiphenotypic population changes through time in a manner that decreases survival at peaks in the oscillations, and increases survival in troughs, providing an emergent damping force.

We prove global stability for the simple cases illustrated above, and conjecture that this stabilizing effect of intraspecific variation can be found in a much wider class of consumer-resource models. In Fig. 3 and the Supplementary Information we consider some interesting cases. For example, in Fig. 3i we show that the result holds for a model where demographic parameters are inherited with some fidelity (i.e., each phenotype preferentially gives rise to offspring of the same type). This feature turns our model into a simple and tractable eco-evolutionary system. We also note that theoretical results derived under the assumption of homogeneous populations can be complicated by the inclusion of intraspecific variation. For example, Fig. 3ii shows *n* predator species coexisting on *k* < *n* prey species—which would be impossible under classic consumer-resource dynamics, even in the presence of strong self-regulation ^8,13^.

**Figure 3:**
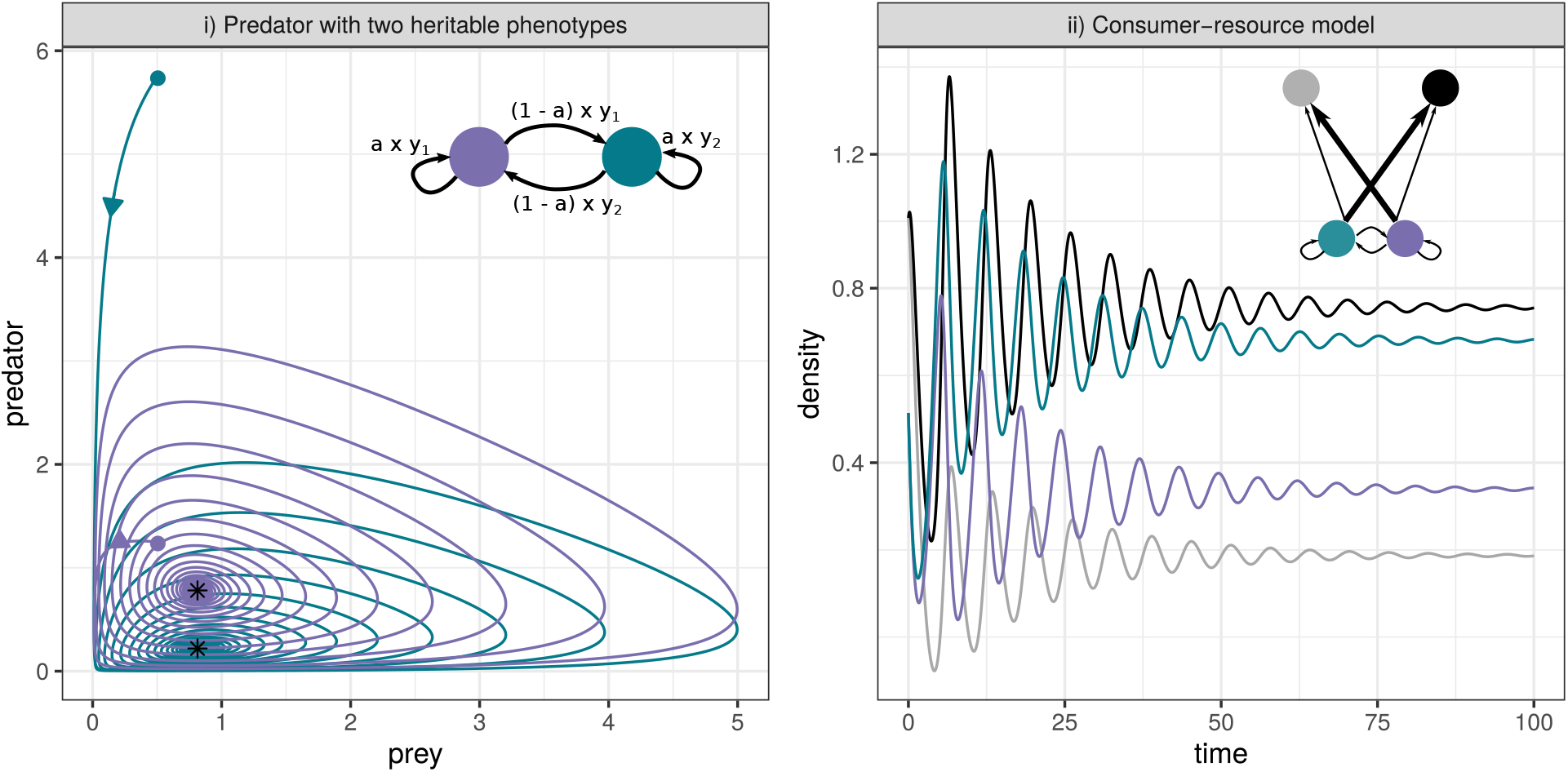
More complex models. i) As in Fig. 1ii, but with heritable predator phenotypes. Instead of pooling reproduction and dividing it equally among phenotypes, each predator phenotype gives rise to offspring of the same phenotype with probability *a*, and of the other phenotype with probability (1 − *a*) (reproduction structure detailed in the inset); dynamics are still stable for this case (here, *δ* = 1/3, *a* = 3/4). ii) A consumer-resource model in which two consumers (grey, black) prey preferentially on different prey phenotypes (colors, structure of the food web in the inset), leading to the stable coexistence of *n* consumers on *k* < *n* prey, contrary to what is found for simple consumer-resource models with homogeneous populations ^8,13^. Parameters are reported in the Supplementary Information.

Overall, these results demonstrate that intraspecific variability can have a substantial impact on the dynamics of ecological systems, qualitatively and quantitatively altering outcomes, and requiring us to amend and extend well-established results. In our models, which are minimal extensions of the classic Lotka-Volterra predator-prey equations to include intraspecific variation, dynamics are always stable despite the absence of any intraspecific competition or other self-regulation, providing an alternative (or complementary) pathway to stable coexistence in diverse ecological systems.

Our work connects with a large and growing body of literature ^18–21^ on the effects of intraspecific variability in ecological systems, providing a simple, striking example of why and how variability matters ^22^. In this context, our results align most strongly with those of Maynard *et al.* ^21^, who analyzed the effect of intraspecific variation on an extension of the replicator-mutator equation, which is closely related to the Lotka-Volterra model ^9^. Moreover, our findings parallel the notable results recently published by de Roos ^23^, who was able to show stabilization of large food webs without widespread self-regulation when accounting for populations structured into juveniles and adults. Foregoing the self-evident truth that not all individuals in a population are created equal has served ecological theory well for over a century, but our work adds to the mounting evidence showing that we need to tackle intraspecific variability to make progress in the discipline.

## Methods

### Lotka-Volterra predator-prey model

Consider a more general parameterization of Eq. 1:

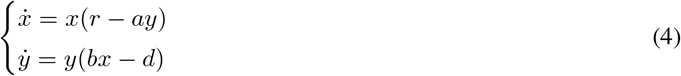

where *r* > 0 models the intrinsic growth rate of the prey, *d* > 0 the mortality rate of the predator, *a* > 0 the attack rate of the predator on the prey and *b* > 0 the positive effect of the prey on the growth of the predator. Besides the trivial equilibrium, we can find the coexistence equilibrium *x*\* = *d/b* > 0, *y*\* = *r/a* > 0. We want to prove that model dynamics form closed orbits around this equilibrium from any positive initial condition. To this end, we write the function:

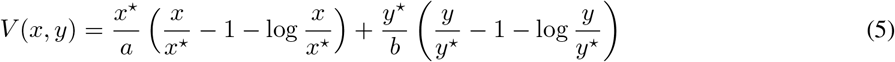

This function is nonnegative whenever *x* > 0 and *y* > 0 (because *x/x*\* > 0, *y/y*\* > 0, and *z* − 1 − log *z* ≥ 0 whenever *z* > 0), and is clearly exactly zero at the coexistence equilibrium. Taking the derivative of *V* (*x, y*) with respect to time we find 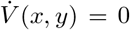, proving that this quantity is conserved through the dynamics, and therefore *V* (*x*(*t*), *y*(*t*)) = *V* (*x*(0), *y*(0)) (i.e., the initial conditions determine the value of *V*). As such, dynamics are constrained to be moving along closed orbits (Fig. 1i). Finally, by computing the flow at each point on this curve, one can prove that dynamics move counterclockwise in the phase plane. The Lotka-Volterra model can be recast as a Hamiltonian system, as found in classical mechanics ^14^.

### Stabilization via self-regulation

The behavior of the model changes qualitatively if the predator or the prey limit their own growth. For example, if we add a term modeling intraspecific competition among the prey, we obtain:

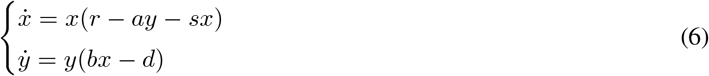

with *s* > 0. The equilibrium becomes *x*\* = *d/b* > 0, *y*\* = *r/a* − *ds/*(*ab*), which is positive whenever all parameters are positive, and *br* < *ds*. If the equilibrium is positive (feasible), we can write the same function *V* (*x, y*) in Eq. 5, which again is positive for all *x, y* > 0, except at the coexistence equilibrium. Differentiating with respect to time, we find:

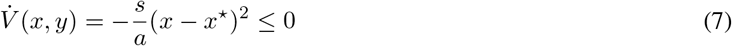

which is negative for any *x* > 0 besides *x* = *x*\*. As such, *V* (*x, y*) is a Lyapunov function (Supplementary Information) for the system, proving the global stability of the coexistence equilibrium.

Note that when 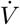 is not strictly negative everywhere but at the equilibrium (as in this case—the derivative is zero whenever the prey is at its equilibrium abundance, regardless of the abundance of the predator) one needs an extra step to prove stability; in the remainder of the Methods we prove nonpositivity of 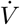, and we provide a more thorough analysis in the Supplementary Information.

The same can be proven when the predator is (or both species are) self-regulating.

### Global stability for the model with two predator phenotypes

We want to show that the coexistence equilibrium for the model in Eq. 2, *x*\* = 1 − *δ*^2^, 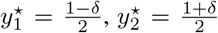 is globally stable for any 0 < *δ* < 1. We consider the function:

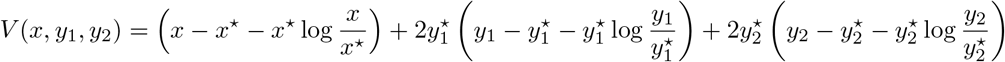

which is nonnegative whenever *x, y*_1_, *y*_2_ > 0 and is zero at the coexistence equilibrium. Differentiating with respect to time, we obtain:

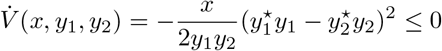

proving the global stability of the coexistence equilibrium.

### Global stability for the model with two prey phenotypes

For the model in Eq. 3, the coexistence equilibrium is given by 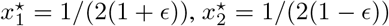, *y*\* = 1/(1 − *ϵ*^2^), which is feasible for any 0 < *ϵ* < 1. We take the function:

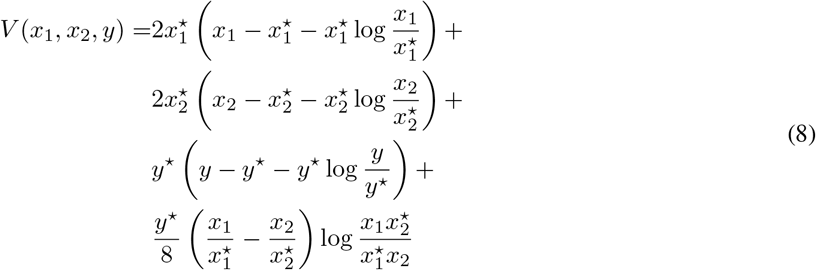

and notice that the function is nonnegative whenever *x*_1_, *x*_2_, *y* > 0, because each of the four terms is nonnegative. The function is also zero at the equilibrium. In the Supplementary Information, we prove that the derivative with respect to time is nonpositive, 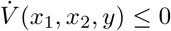, thereby ensuring the global stability of the equilibrium.

### Multiple predator phenotypes

One can extend the model in Eq. 2 to the case of an arbitrary number of predator phenotypes, *n*:

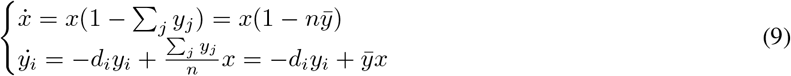

where we assume that *d*_*i*_ > 0, ∑_*i*_ *d*_*i*_ = *n*, and 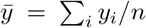. At the coexistence equilibrium, *x*\* = *H* and 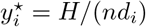, where 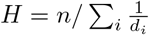, the harmonic mean of the predator mortality rates. Therefore, 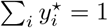. We write the Lyapunov function:

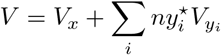

with 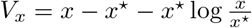 and 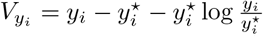. The function is nonnegative when all abundances are positive, and zero at the coexistence equilibrium. Differentiating with respect to time, we obtain:

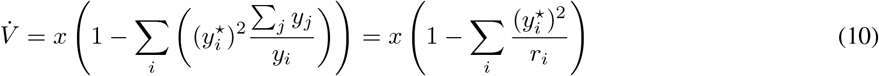

where we have defined *r*_*i*_ = *y*_*i*_/ ∑_*j*_ *y*_*j*_. Radon’s inequality states that, for *α*_*i*_, *β*_*i*_ > 0, 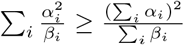. Choosing 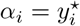 and *β*_*i*_ = *r*_*i*_, we find 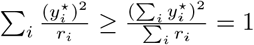, proving that 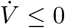, which, as we show in the Supplementary Information, guarantees the global stability of the coexistence equilibrium.

Note that in Eq. 9 reproduction is partitioned equally among the *n* phenotypes. However, we can exploit these equations to model dynamics for the case of a predator characterized by an arbitrary distribution of mortality rates. Suppose that, at birth, a predator is assigned a mortality rate *d* with probability *P* (*d*). We can approximate any distribution *P* (*d*): a) take *n* to be large, b) for each *d*, assign to *k* phenotypes mortality rate *d*, such that *k/n* ≈ *P* (*d*). Because the dynamics of all the phenotypes with the same mortality rate will rapidly synchronize, the two models are equivalent. By taking *n* to be sufficiently large, we can approximate any *P* (*d*) with arbitrary precision.

## Acknowledgements

P. Lemos-Costa, A. Skwara and P. Lechon for discussion.

## Code availability

Code reproducing all figures is available at github.com/StefanoAllesina/LV_intra.

## Author contributions

SA and CAS conceived the models; SA, ZRM and CAS performed the mathematical analysis; SA wrote the paper and the code. All authors discussed the results and edited the paper.

## Supplementary Information

### Mathematical preliminaries: stability and Lyapunov functions

Provided with a dynamical system defined by a set of equations 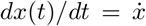, and an equilibrium *x*\* such that 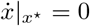, we want to determine the stability of the equilibrium. The equilibrium *x*\* is stable if for any choice of *ϵ* > 0 we can find a positive constant *δ* such that if ||*x*(0) − *x*\*|| < *δ*, then ||*x*(*t*) − *x*\*|| < *ϵ* for every *t* ≥ 0. If for *t* → ∞ *x*(*t*) → *x*\* the equilibrium is asymptotically stable. For a stable equilibrium, we can identify a region of attraction, i.e., the set of initial conditions asymptotically leading to *x*\*.

In 1892, Aleksandr Lyapunov ^24^ proposed a general method to determine the stability of an equilibrium for a given region of initial conditions. For ecological models with *n* populations (or phenotypes), we are interested in the region of the *n*-dimensional space in which all quantities are positive (i.e., the positive orthant of ℝ^*n*^, denoted 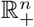). Lyapunov’s method (called the “direct method”) is very simple: typically, we cannot write the solution for the evolution of the dynamical system of interest in time starting from arbitrary initial conditions; however, if we a) define a space *D*, containing the equilibrium *x*\*, and b) identify a function of the variables constituting the system such that:

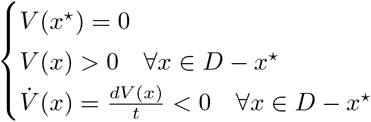

then *x*\* is stable. Moreover, if 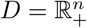 and *V* (*x*) is radially unbounded (i.e., *V* (*x*) → ∞ when *x* → *∂D*) then *x*\* is globally asymptotically stable in 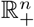. This condition on *V* guarantees that the trajectory does not escape from *D* as *t* → ∞.

Intuitively, we can think of *V* (*x*) as a function summarizing the state of the system; because *V* (*x*) > 0 everywhere besides at the equilibrium, and is decreasing in time, then it follows that the system will eventually reach the equilibrium.

Thanks to LaSalle’s invariance principle, the condition can be relaxed to 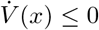 for many systems of interest. In such cases, if the equilibrium is the only trajectory contained in the set defined by 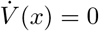, stability is guaranteed.

While Lyapunov’s direct method allows one to prove stability in a straightforward way if provided with a function *V* (*x*), the crux of the approach is the identification of a suitable function out of infinitely many choices. A number of algorithms to solve this issue have been proposed ^25^, but this task can be very daunting: as Strogatz ^26^ put it, “divine inspiration is usually required”.

### The Generalized Lotka-Volterra model

The Generalized Lotka-Volterra model is arguably the simplest set of equations describing the population dynamics of a set of interacting species, and has served as the backbone for the development of ecological and epidemiological models for over a century.

In its predator-prey incarnation, it was first proposed in 1920 by polymath and peripatetic scientist Alfred J. Lotka ^1^, who devised a number of chemical/ecological models. Lotka clearly had an ecological system in mind, considering *“a plant species […] deriving its nourishment from a source presented in such large excess that the mass of the source may be considered constant during the period of time with which we are concerned.”* and *“a herbivorous animal species”* feeding on the first species. In his original paper, Lotka Taylor-expanded the dynamics of a perturbation around the coexistence equilibrium and showed that what we now call the “community matrix” associated with this equilibrium has purely imaginary eigenvalues—indicating that the equilibrium is neutrally stable. He recognized that in such case the equations describing the linearized dynamics of a small perturbation took the form of a Fourier’s series, therefore concluding that the system would oscillate indefinitely. The cyclic behavior of the solution ignited Lotka’s imagination—he saw the potential for such equations to describe the “rhythm of Nature” dear to the Victorians, and in fact he included in the paper a long quote from the Victorian polymath Herbert Spencer. Lotka continued to study this model, and in a subsequent paper ^2^, he was able to show that indeed predators and prey cycle around the neutral equilibrium in a closed orbit, and to identify the constant of motion determining the trajectory.

Intriguingly, the same exact model and solution were independently proposed in 1926 (in a long, mathematically sophisticated study ^4^ and in a short note in *Nature* ^3^, devoid of equations) by Italian mathematician Vito Volterra, who had been inspired by the work of his son-in-law, fish-biologist Umberto D’Ancona. D’Ancona had noticed that the closure of fisheries in the Adriatic sea due to World War I had benefited predatory fish, but not herbivorous ones ^5^, and asked “papà” ^32^ whether this pattern could be explained mathematically. Volterra, then aged 66, was a prominent mathematician, patriot, and Senator of the Kingdom of Italy, and was about to be expelled (in 1931) from Italian academia for refusing to swear his allegiance to the fascist regime, which he vehemently opposed. Volterra considered not only predator-prey interactions, but also more general cases including, for example, species competing for shared resources. He also extended the model to an arbitrary number of species.

Lotka wrote to *Nature* in 1927 to claim priority ^33^, and concluded that *“It would be gratifying if Prof. Volterra’s publication should direct attention to a field and method of inquiry which apparently has hitherto passed almost unnoticed.”* Volterra graciously recognized Lotka’s priority ^34^, and agreed that *“these studies and these methods of research deserve to receive greater attention from scholars, and should give rise to important applications.”* The research program initiated by Lotka and Volterra became hugely successful indeed: *“these studies and these methods”* are now part of the canonical body of work taught to every ecologist around the world, and have been extended and analyzed in countless studies. Moreover, their equations arise in a variety of fields including laser dynamics ^35,36^ and plasma physics, communication networks ^37^, game theory ^9^ and epidemiology—and a much wider class of models can be recast as a larger-dimensional Lotka-Volterra model in which dynamics are constrained to a lower-dimensional manifold ^39–41^.

For a system of *n* interacting species, the Generalized Lotka-Volterra model (GLV) can be written as:

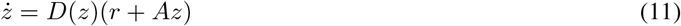

where *D*(*z*) is a diagonal matrix with the density of all species at time *t*, *z*(*t*), on the diagonal, *r* is a vector of intrinsic growth rates (for producers) or mortality rates (for consumers), and *A* is a matrix representing species interactions (with the diagonal representing “self-interactions”—for example intraspecific competition). If a feasible (i.e., positive) equilibrium exists, it is unique (when matrix *A* has full rank) and is computed as *z*\* = −*A*^−1^*r*. The existence of a feasible equilibrium is necessary (but not sufficient) for coexistence 9. Whenever a feasible equilibrium exists, and appears to be globally stable, one can attempt applying Lyapunov’s direct method via the candidate Lyapunov function proposed by Goh 42, which reduces to the function in Eq. 5 for the predator-prey case. We examine the derivation of this function in some detail as it serves as a jumping board for our own analysis.

To produce a Lyapunov function, we need to start with a function that is zero at equilibrium and positive anywhere else in the region of interest. Goh’s function exploits the fact that, for any *x* > 0, *x* − 1 − log *x* ≥ 0, with equality attained at 1. Take *x* to be 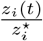 (which is positive whenever the equilibrium is feasible and *z*_*i*_ (*t*) > 0), then 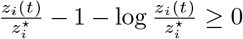, with equality attained at equilibrium. Multiplying by 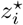 we obtain 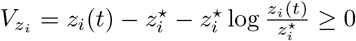. Our candidate Lyapunov function is therefore:

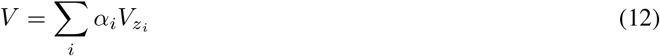

which is positive for any choice of *α*_*i*_ > 0 whenever *z*_*i*_ > 0 for all *i* and *z* ≠ *z*\*; the function is zero at equilibrium. Thus, if we can identify positive constants *α*_*i*_ such that 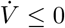, we have proven the stability of the equilibrium. For the GLV model, we can simplify the equations by noting that *r* = −*Az*\*, and therefore 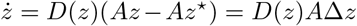 (where Δ*z* is the vector of deviations from equilibrium, *z* − *z*\*). Differentiating Eq. 12 with respect to time, we find:

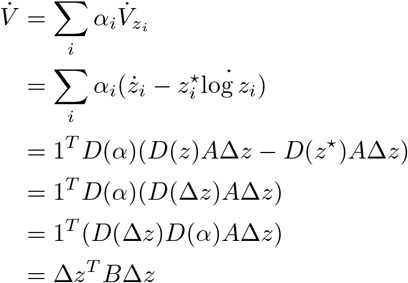

where *D*(*α*) is a diagonal matrix with the *α*_*i*_ on the diagonal, *B* = *D*(*α*)*A*, 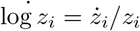, and we have exploited the fact that diagonal matrices commute. For stability, we need to prove that Δ_*z*_^*T*^ *B*Δ_*z*_ ≤ 0. If a symmetric matrix *C* has the property that *x*^*T*^ *C*_*x*_ ≤ 0 for every *x* ∈ ℝ^*n*^, it is called negative semi-definite. Clearly, only the symmetric part of *B*, (*B* + *B*^*T*^)/2 matters for the sum (because the skew-symmetric part cancels). As such, if we can find a set of positive constants *α*, such that (*B* + *B*^*T*^)/2 = (*D*(*α*)*A* + *A*^*T*^ *D*(*α*))/2 is negative semi-definite, we have proven the stability of the coexistence equilibrium. Note that this is a sufficient, but not necessary condition.

### Stability for a model with multiple predator phenotypes

In the Main Text, we set all parameters besides those pertaining to phenotypic differences to one, to highlight a minimal setting in which our results hold. Naturally, these results can be extended to more realistic cases, as found for the predator-prey model (cfr. Eq. 1 and Eq. 4). For example, we can introduce demographic parameters to extend the model with *n* predator phenotypes (Eq. 9):

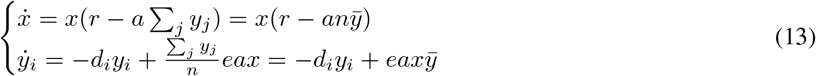

where we have set a growth rate for the prey (*r* > 0), an attack rate for the predator phenotypes (*a* > 0) and an efficiency of transformation (*e* > 0). We assume that *d*_*i*_ > 0∀*i* and, without loss of generality, that ∑_*i*_ *d*_*i*_ = *n* (if this is not the case, simply rescale time to recover this parameterization).

From the equation for the prey dynamics, we have that, unless *x* = 0, at equilibrium 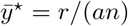. From this, we can find 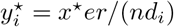; by summing across 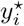 we can solve *x*\* = *n*/(*ea* ∑_*i*_(1/*d*_*i*_)) = *H/ea*, where *H* is the harmonic mean of the predator phenotypes’ mortalities. In summary, at the feasible coexistence equilibrium we have:

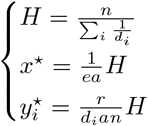

We want to prove the stability of this equilibrium. As before, we define the nonnegative functions 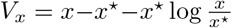 and 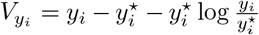 and attempt to find a suitable Lyapunov function with the same form as that proposed by Goh ^42^:

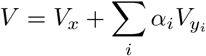

where *a*_*i*_ > 0∀*i*. A convenient choice is 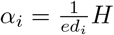, leading to:

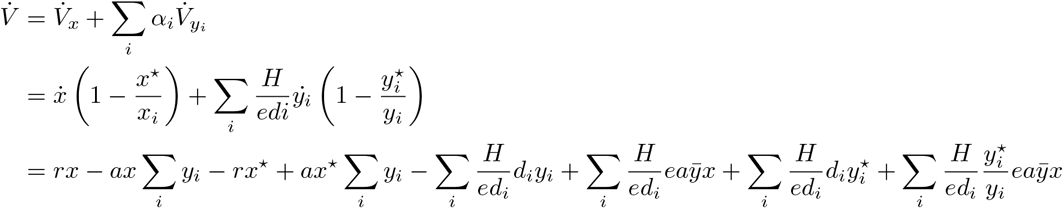

We can cancel three pairs of terms:

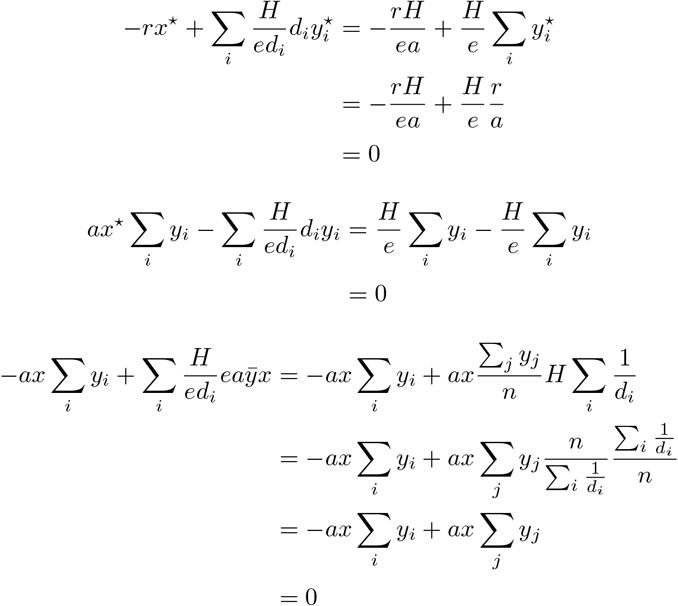

We are left with:

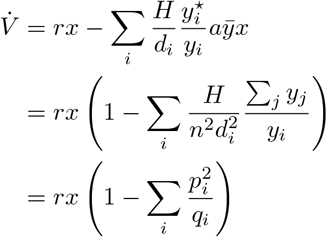

where we have defined 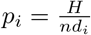 and 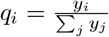. Applying Radon’s inequality, we find:

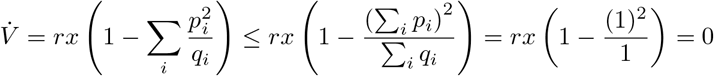

proving that *V* does not increase in time. Given that *V* is unbounded towards the boundaries of the positive orthant, to conclude that the equilibrium is stable it is sufficient to show that the equilibrium is the only trajectory contained in 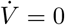. Any such trajectory will satisfy 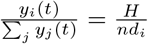. This implies that 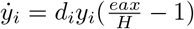. Because the ratio 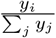 is constant, it follows that:

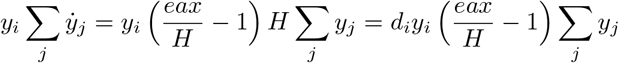

As *d*_*i*_ ≠ 1 and *H* ≠ 1 we must have *eax* = *H* and thus 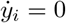. This further imposes that 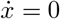, concluding the proof.

### Stability for a model with two prey phenotypes

Proving global stability for the case of a predator and two prey phenotypes coupled by reproduction (Eq. 3) is more complicated, because we cannot find suitable constants for the (relatively) simple candidate Lyapunov function in Eq. 12. We therefore use the modified Lyapunov function (Eq. 8):

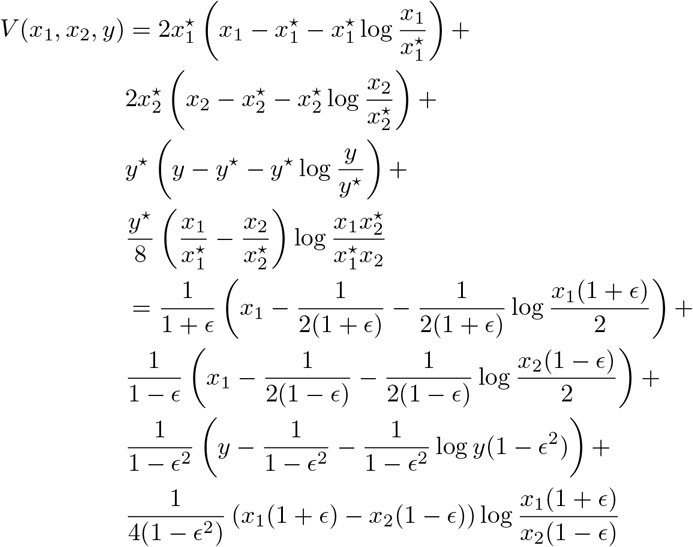

As discussed in the Main Text, this function is nonnegative for all positive abundances, and zero at equilibrium. In order to prove that *V* is indeed a Lyapunov function, it remains to show that 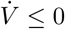.

In the calculations that follow, it will be convenient to use the new variables *z*_1_ = (1 + *ϵ*)*x*_1_ and *z*_2_ = (1 − *ϵ*)*x*_2_. Making these substitutions and differentiating with respect to time, we obtain:

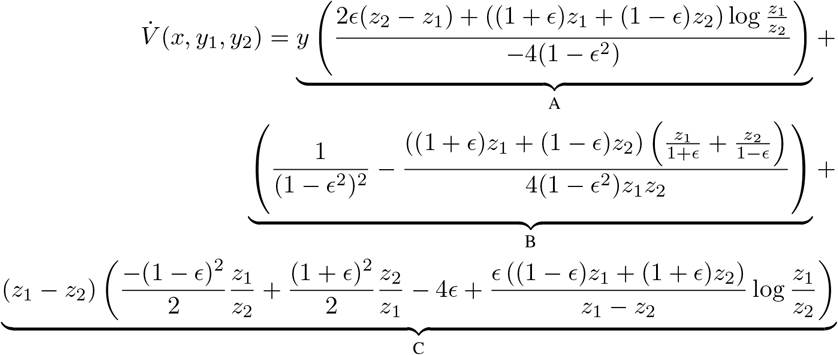

We will prove that all of the three labeled terms, A, B, and, C, are independently nonpositive. We rely on the positivity of the variables *z*_1_, *z*_2_, and *y* throughout.

### Term A

The factor 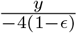 is strictly negative, so A is nonpositive if and only if

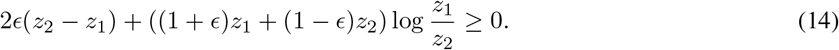

Re-grouping terms, Eq. 14 becomes

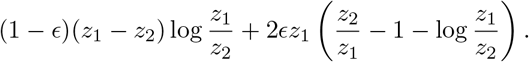

The left-hand term is nonnegative because *z*_1_ − *z*_2_ and 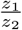 must share the same sign. The right-hand term is also nonnegative, due to the inequality *x* − 1 − log *x* ≥ 0 for *x* > 0. Consequently, term A is always nonpositive through the dynamics.

### Term B

Term B is nonpositive if the following inequality holds:

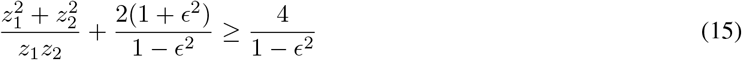

The first term on the left-hand side is bounded from below by 2 (this is a consequence of (*z*_1_ − *z*_2_)^2^ > 0). So we have

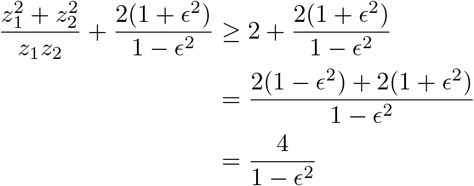

which establishes Eq. 15.

### Term C

We prove that term C is nonnegative by considering two cases: *z*_2_ > *z*_1_ and *z*_1_ > *z*_2_ (when *z*_1_ = *z*_2_, we have C = 0). In each case, the sign of *z*_1_ − *z*_2_ is fixed, so we concentrate on bounding the terms in parentheses, which we denote by *s*. First, when *z*_2_ > *z*_1_, C is nonnegative if

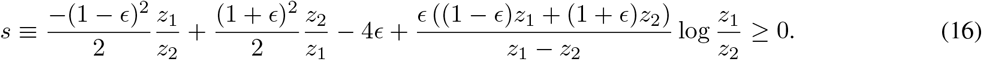

We can easily lower-bound the first two terms in *s* using the assumption *z*_2_ > *z*_1_. The result is

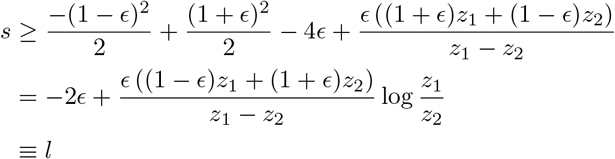

Now we employ a bound on the logarithm which holds for 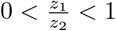:

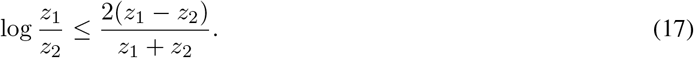

This inequality is easily derived from the fact that 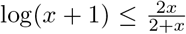 for −1 < *x* < 0 (see, e.g., ^43,44^). Because the logarithmic term in Eq. 16 is multiplied by a negative coefficient, Eq. 17 yields a lower-bound:

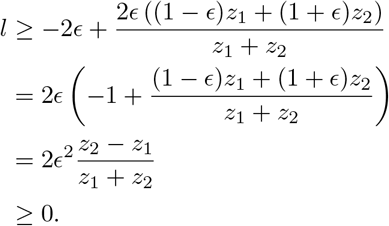

This series of bounds establishes that Eq. 16 holds, and so C is nonpositive when *z*_2_ > *z*_1_.

Finally, we consider the case *z*_1_ > *z*_2_. Now the factor *z*_1_ − *z*_2_ is positive, so C is nonpositive if

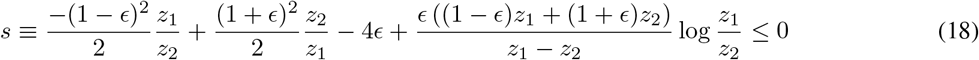

(the reverse of Eq. 16). We note that

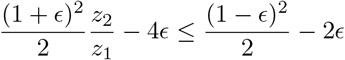

so we have

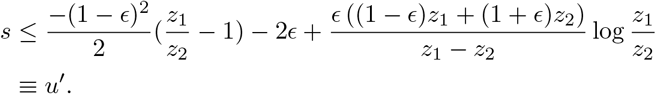

This expression can be simplified considerably by introducing the new variable 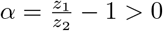:

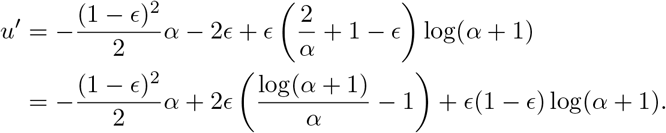

Here we use another logarithmic bound,

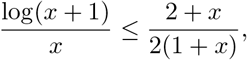

which holds for *x* > 0 (^44^). With this inequality, a second upper bound on *s* is

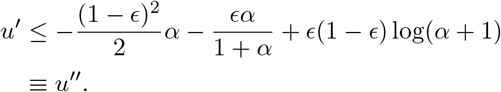

At last, we employ one trivial 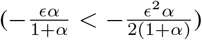 and one non-trivial inequality 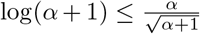; see ^43^) to obtain a final upper bound:

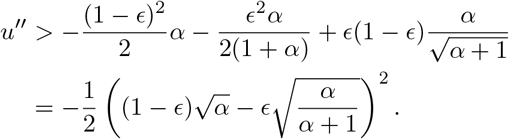

This last expression is, of course, nonpositive, showing that *s* ≤ 0 (Eq. 18). Consequently, C is nonpositive when *z*_1_ > *z*_2_, as well as when *z*_2_ > *z*_1_ (above). We conclude that C is nonpositive for any choice of the variables.

Having shown that each of A, B, and C is nonpositive, it must be the case that *V* decreases through the dynamics. Thus, *V* is a Lyapunov function for the model with two prey phenotypes (Eq. 3 in the Main Text), and the equilibrium for this model is globally stable. To conclude this, we rely on LaSalle’s invariance principle: Due to the interplay between the linear and log terms, *V* is (positively) unbounded towards the boundary of the positive orthant. Thus, we only need to show that the only trajectory contained in 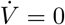 is the equilibrium. A close examination of the section above shows that *C* is strictly negative for any *z*_1_ ≠ *z*_2_, so any trajectory {(*x*_1_(*t*), *x*_2_(*t*), *y*(*t*))} contained on the set 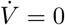 satisfies 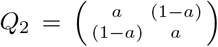. Using Eq. 3 in the Main Text, we conclude that 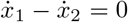, and consequently *x*_1_(*t*)*, x*_2_(*t*) and *y*(*t*) are constant. Hence, the only trajectory contained in 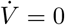 is the equilibrium and the claim follows.

### Generalizing Lotka-Volterra

All the models presented here and in the Main Text can be seen as a further generalization of the GLV equations. We can rewrite the GLV model as:

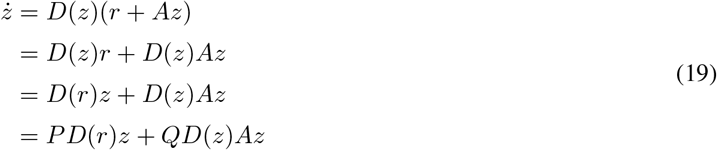

where matrices *P* and *Q* are the identity matrix. The models presented here are of this form, with the difference that matrices *P* or *Q* are block-structured, rather than being the identity matrix. In particular, the matrix *P* describes how the reproduction of the producer phenotypes is distributed among the phenotypes forming a population, and the matrix *Q* how the reproduction of the consumer phenotypes is apportioned.

For our models, we have *n* populations, and each population *i* is composed of a certain number of phenotypes, *k*_*i*_, connected by reproduction. As such, defining *m* = Σ_i_ *k*_i_, we have that the vector *r* is of length *m*, and contains the phenotype-specific growth/mortality rates; similarly, matrix *A* is of size *m* × *m*, with each coefficient *A*_*jl*_ representing the effect of phenotype *l* on the growth of phenotype *j*. If we order the equations such that all phenotypes belonging to a population are arranged consecutively, we can define *P* and *Q* as block-diagonal matrices, with blocks *P*_1_, …, *P*_*n*_ on the diagonal of *P* (of size *k*_1_, …, *k*_*n*_, respectively), and similarly *Q*_1_, …, *Q*_*n*_ for matrix *Q*. Producers reproduce in the linear part of the equation, and consumers in the quadratic part; therefore we take *P*_*i*_ to be a nonnegative, irreducible, and column-stochastic *k*_*i*_ × *k*_*i*_ matrix if population *i* represents a producer, and 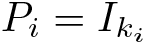 (the identity matrix of size *k*_*i*_) otherwise; conversely, 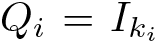 for producers, and is a column-stochastic, nonnegative, irreducible matrix for consumers. Irreducibility here models the concept of a biological species: through reproduction we can go from any phenotype to any other within a population. For the matrix *A* we have the form 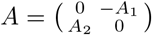, where *A*_1_ and *A*_2_ are of suitable size and nonnegative. Note that if we want to introduce self-effects, or species that are both consumers and consumed, we need to complicate the equations slightly, because only reproduction (but not the competition/predation) should be apportioned by *Q*. A possibility is to split *A* into *A* = *A*^(+)^ + *A*^(−)^, where *A*^(+)^ stores the coefficients pertaining to the reproduction of the consumer species, and *A*^(−)^ the terms representing competition or predation. In this more complex case, we have:

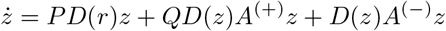

For simplicity, here we keep to the case of species cleanly divided into predators and prey, and not experiencing intraspecific competition. For example, Eq. 2 of the Main Text (two predator phenotypes) can be recast in this form by defining:

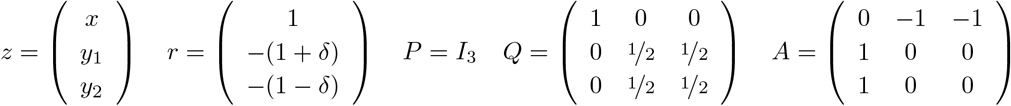

Similarly, Eq. 3 can be rewritten as:

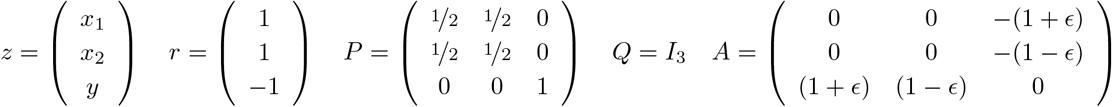

This generalization encompasses all the models presented here, and is clearly related to the replicator-mutator equation found in evolutionary game theory ^21,45^. The model appears to be quite difficult to analyze in the general case—for example, even writing the equilibrium analytically can be easily accomplished only for contrived or small-dimensional parameterizations.

### Simulations

Because Eq. 19 is difficult to analyze in the general form with *P* and *Q* being column-stochastic, block-structured matrices, in this section we present simulations describing two interesting cases, reported in Fig. 3 of the Main Text, as well as a more complex case in which both consumers and resources display intraspecific variation.

#### An eco-evolutionary model

In the models above, newborns are assigned a certain death rate (or attack rate) at birth, according to some random distribution. A more biologically-interesting case is one in which the lifespan is (or other parameters are) inherited with some fidelity from the parents ^21^. This feature turns our model into a simple eco-evolutionary system, in which for example “mutations” in asexually reproducing populations connect phenotypes belonging to the same population. Take for example a model with two predator phenotypes and one prey:

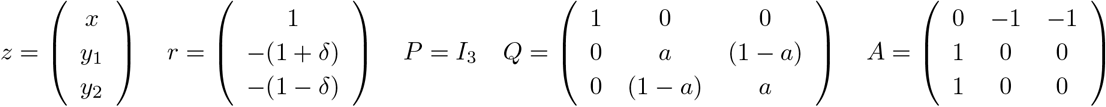

which is equivalent to Eq. 2, but in the more general case in which 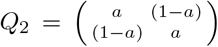, rather than having all coefficients equal to 1/2. Here *a* represents the probability with which a phenotype is inherited, and 1 − *α* the probability that a newborn of the other phenotype is produced. Clearly, one can generalize this further, by allowing for any nonnegative, irreducible, and column-stochastic *Q*_2_, and/or a different number of populations. Fig. 3i shows that this generalization leads to the stabilization when *δ* = 1/3 and *α* = 3/4.

#### A consumer-resource model

MacArthur’s celebrated consumer-resource framework can be rewritten as a GLV model ^8,13^. We can therefore use Eq. 19 to introduce intraspecific-variation in the consumer(s) or producer(s), generalizing the model. Coexistence of *n* consumers on *k* < *n* resources (or more generally *n* populations and *k* < *n* regulating factors ^49^) is an impossibility in the classic formulation of the model ^50^, as already outlined by Volterra ^4^. The argument for “competitive exclusion” was extended (and hotly debated) in subsequent papers ^49–51^. In our model, it is certainly possible to have stable dynamics for a smaller number of prey than predators, for example when consumers prefer different prey phenotypes. Take:

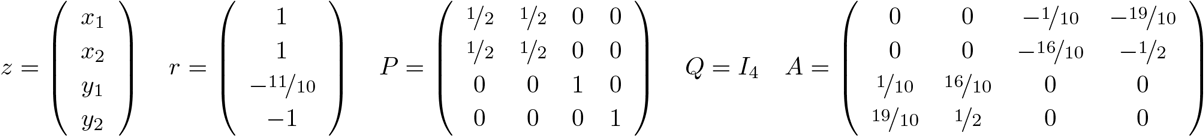

Simulations using these parameters (Fig. 3ii) yield stable coexistence. The results are therefore similar to those found by Haigh and Maynard Smith ^51^ for a structured prey population composed of juveniles and adults.

Larger communities can be stabilized by intraspecific variation. For example, taking Eq. 19 with parameters:

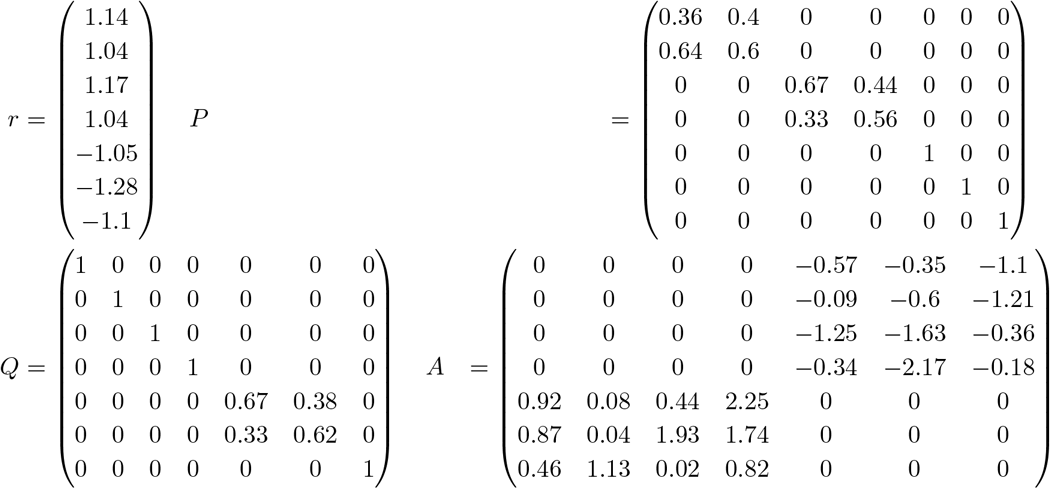

i.e., a model with two producers (each composed of two phenotypes) and two consumers (one with two phenotypes and the other with a single phenotype), dynamics converge to the feasible equilibrium (Fig. 4).

**Figure 4:**
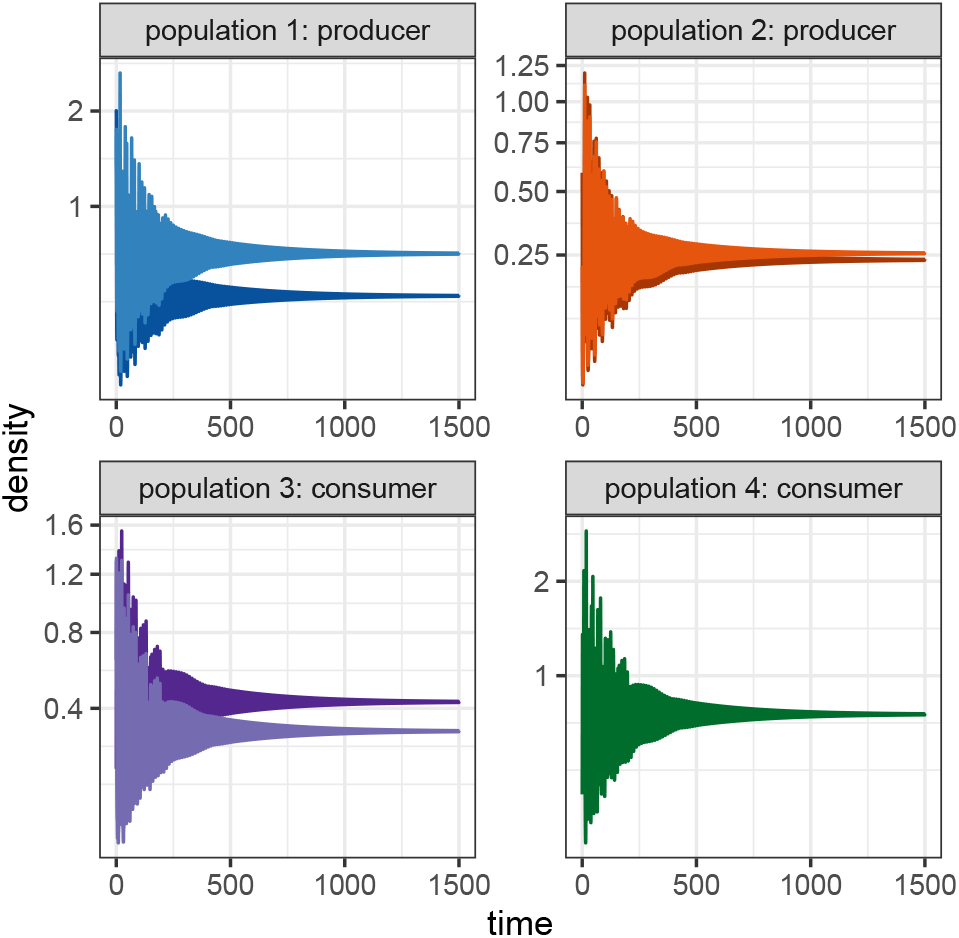
A larger community. Stable dynamics for a model with two producers and two consumers.

